# An Optimal Control Approach for Enhancing Natural Killer Cells’ Secretion of Cytolytic Molecules

**DOI:** 10.1101/2020.08.05.238691

**Authors:** Sahak Z. Makaryan, Stacey D. Finley

## Abstract

Natural killer (NK) cells are immune effector cells that can detect and lyse cancer cells. However, NK cell exhaustion, a phenotype characterized by reduced secretion of cytolytic models upon serial stimulation, limits the NK cell’s ability to lyse cells. In this work, we investigated *in silico* strategies that counteract the NK cell’s reduced secretion of cytolytic molecules. To accomplish this goal, we constructed a mathematical model that describes the dynamics of the cytolytic molecules granzyme B (GZMB) and perforin-1 (PRF1) and calibrated the model predictions to published, experimental data using a Bayesian parameter estimation approach. We applied an information-theoretic approach to perform a global sensitivity analysis, from which we found the suppression of phosphatase activity maximizes the secretion of GZMB and PRF1. However, simply reducing the phosphatase activity is shown to deplete the cell’s intracellular pools of GZMB and PRF1. Thus, we added a synthetic Notch (synNotch) signaling circuit to our baseline model as a method for controlling the secretion of GZMB and PRF1 by inhibiting phosphatase activity and increasing production of GZMB and PRF1. We found the optimal synNotch system depends on the frequency of NK cell stimulation. For only a few rounds of stimulation, the model predicts inhibition of phosphatase activity leads to more secreted GZMB and PRF1; however, for many rounds of stimulation, the model reveals that increasing production of the cytolytic molecules is the optimal strategy. In total, we developed a mathematical framework that provides actionable insight into engineering robust NK cells for clinical applications.

## 1 INTRODUCTION

Natural killer (NK) cells are innate immune effector cells that protect the host from diseased cells such as virally-infected cells and cancer cells^1,2^. In particular, when NK cells engage with these target cells, NK cell stimulatory receptors become activated and mediate killing of the diseased cells. One mechanism for target cell-killing is through the secretion of the cytolytic molecules granzyme B (GZMB) and perforin-1 (PRF1)^3–6^. Secretion of these factors is termed “degranulation”. Specifically, PRF1 mediates the formation of pores on the target cell membrane, enabling GZMB to infiltrate and induce apoptosis. Although the secretion of cytolytic molecules is mediated by multiple NK cell receptor signaling pathways^7^, including – but not limited to – the natural cytotoxicity receptors (e.g., NKp46), 2B4 (CD244) and DNAM-1 (CD226), the CD16 and NKG2D receptors are two of the most studied. In fact, a significant majority of NK cells *in vivo* are CD16-positive. Specifically, CD16 is an Fcγ receptor found on the surface of NK cells^7–10^, which binds to the constant region of immunoglobulin G (IgG) antibodies. Due to its affinity for antibodies, CD16 is necessarily required for antibody-dependent cell-mediated cytotoxicity (ADCC), a mechanism for lysing target cells through antibodies. This feature of the CD16 receptor has been integral for designing bi- and tri-specific killer engagers (BiKEs and TriKEs)^11,12^, which are engineered antibodies that traffic NK cells to target cells for cell killing. NKG2D belongs to the CD94/NKG2 family of receptors and has been found on NK cells as well as T cells^13–15^. Unlike CD16’s ubiquity in ADCC, NKG2D is specific as it recognizes and binds to induced-self antigens (e.g., MHC class I polypeptide-related sequence A (MICA)) on the surface of cells. These antigens communicate to NK cells that the diseased cell should be lysed. This implicates NKG2D in the elimination of diseased cells, including cancer cells. Excitingly, NKG2D serves as a focal point for many lines of research in targeted therapies^15–18^ due to its affinity for tumor-associated antigens.

While CD16 and NKG2D are activated under different biological scenarios, they activate a similar set of downstream signaling molecules^7–10,19^ that mediate the secretion of GZMB and PRF1. Upon binding to their cognate ligands, antibodies, or antigens, CD16 and NKG2D promote activation of the Src family kinases (SFK) through the intracellular adaptor molecules CD3ζ and DAP10, respectively. The activation of SFKs leads to the phosphorylation of downstream signaling species PLCγ, Vav, SLP76, Akt, and Erk, as well as phosphatases SHP and SHIP^8–10,20,21^. The phosphatases inhibit the activation of the signaling intermediates, and thus prevent cell activation. The activation of Vav and SLP76 are critical for actin remodeling and formation of the immunological synapse^4,22,23^ between the NK cell and the target cell. Moreover, phosphorylation of Akt and Erk have been correlated with cell survival and proliferation, respectively^7,24–26^. Studies have shown^3,4,8,9,22,27–29^ a strong correlation between phosphorylation of the signaling molecules Vav and PLCγ, secretion of GZMB and PRF1, and NK cell cytotoxicity, suggesting the activation of these molecules precedes target cell death.

Several lines of research^30–32^ have reported NK cell exhaustion as a consequence of over-stimulation of NK cell receptors. NK cell exhaustion is a phenotype characterized by a decrease in NK cell effector functions (e.g., GZMB/PRF1 secretion)^33^ even with more receptor stimulation. In addition, NK cell exhaustion is correlated with a decrease in the density of stimulatory receptors^30^. Sanchez-Correa, et. al.^34^ found a down-regulation of DNAM-1 in primary NK cells exposed to DNAM-1 ligands CD112 and CD155 expressed by leukemic cells *in vitro*, leading to a dampened immune response upon subsequent stimulation of DNAM-1. Paul and colleagues^35^ observed a reduced quantity of NKG2D-positive NK cells inside the tumor micro-environment (TME) of B16F10-induced melanoma in C57BL/6 mice as opposed to NK cells in the periphery of the tumor site. Moreover, Paul, et. al.^35^ discovered that NK cells within the TME had reduced levels of PRF1 mRNA, as well as lower levels of the cytokine IFNγ and CD107 (a marker of degranulation), compared to NK cells in the periphery, suggesting the TME can co-opt NK cells and promote the exhausted phenotype. In the clinical setting, a decrease in the quantity of NK cell stimulatory receptors and effector molecules have been correlated with poor prognoses for pancreatic, gastric and colorectal cancer patients^36,37^. It follows that the efficacy of NK cell-based adoptive cell therapies will be limited unless NK cells can be modified to overcome the effects of exhaustion. Unfortunately, there appear to be many mechanisms that induce NK cell exhaustion^33^, implying that a single approach may not work to prevent exhaustion. In addition, given the extensive cascade of signaling reactions that mediate NK cell degranulation, and the complex interactions that influence NK cell exhaustion, it is not clear which strategies can be combined to reduce exhaustion or prevent it altogether.

Instead of preventing NK cell exhaustion, we could potentially implement strategies that promote its opposite effect; that is, the continuous secretion of cytolytic molecules. This approach may counterbalance the effects of exhaustion, causing the cell to be more robust to stimulation and thereby allowing the NK cell to kill more target cells. Still, it is not clear how to optimally achieve this objective due to the nonlinearities in cell signaling and activation. Excitingly, mathematical models are useful for providing quantitative insight into complex biological processes, including intracellular signaling leading to immune cell activation^38^. For example, mathematical models of NK cell signaling have contributed to our understanding of NK cell activation: the identification of Vav as the signaling molecule where stimulatory and inhibitory signals integrate to determine activation^29^; the explanation of how weak-affinity stimulatory receptors can counterintuitively inhibit NK cell activation in a non-monotonic manner^39^; the revelation that NK cell signaling occurs at a faster timescale than tumor cell-killing and how modifying antibody concentrations can bridge the two processes closer in time^40^; and the elucidation of strategies that increase the likelihood of NK cell activation^19^ by amplifying the amount of phosphorylated signaling intermediates. Indeed, mathematical models can be applied to address a diversity of research questions, especially in determining optimal strategies, which can save experimental researchers time and resources.

In this work, we have applied a mathematical model to investigate strategies that counteract NK cell exhaustion. We first modified our previous model of NK cell signaling^19^ to include GZMB and PRF1 secretion. The model was first calibrated and validated using published data^27^. We then performed a global sensitivity analysis, revealing that activation of a particular phosphatase strongly influences NK cell secretion of cytolytic molecules. With that information in hand, we simulated the effects of reducing the phosphatase’s impact, alone and in combination with increasing production of cytolytic molecules, for different rounds of stimulation. We found that the optimal strategy for maximizing secretion of cytolytic molecules depends on how many times the NK cells are stimulated: for fewer rounds of stimulation, inhibiting phosphatase activity leads to more secreted GZMB and PRF1; in contrast, for many rounds of stimulation, the production of the cytolytic molecules becomes essential. In conclusion, we constructed a mathematical framework describing the effector function of NK cells, which can aid researchers interested in engineering robust NK cells for clinical applications.

## 2 METHODS

### Construction of the NK cell degranulation model

We adopted the mathematical model of NK cell signaling from our previously published work^19^, which includes a system of nonlinear ordinary differential equations (ODEs) that describes the dynamics of receptors CD16, NKG2D, and 2B4, as well as their downstream signaling intermediates. Here, we focus on CD16 and NKG2D signaling, given extensive experimental data quantifying their roles in promoting cytolytic molecule degranulation. Furthermore, we expanded the model by incorporating the dynamics of GZMB and PRF1 to study and simulate strategies that optimize their secretion. This baseline model is provided in Supplementary File S1 and the list of model species, reactions and parameters are provided in Supplementary File S2. A simplified representation of the baseline model is shown in **Figure 1**, where connections between species represent reaction velocities and are governed by established Michaelis-Menten kinetics.

**Figure 1.**
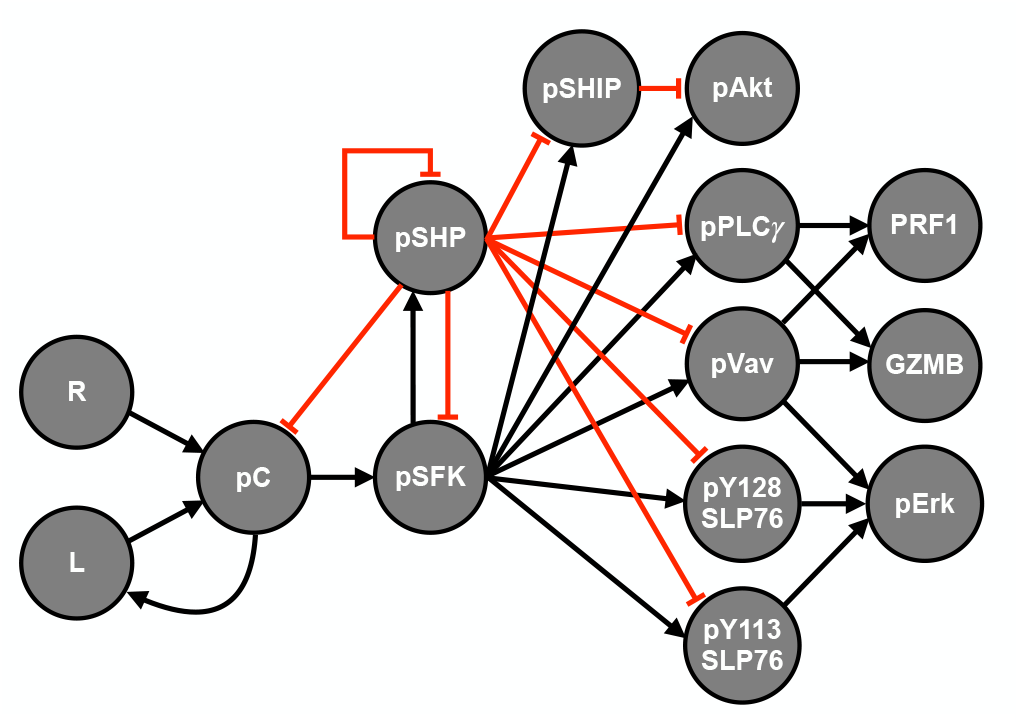
Natural killer cell signaling model schematic. Signal propagation flows from left to right, starting with the interaction between the ligand (L) with the receptor (R) that leads to the activation (phosphorylation) of the receptor-ligand complex (pC). Then, this complex can promote phosphorylation of the Src family kinases (pSFK) which activates the signaling intermediates as well as the inhibitory phosphatases (SHP, SHIP). Finally, the synergistic activation of pVav and pPLCγ influences the release of granzyme B (GZMB) and perforin-1 (PRF1). Arrows indicate stimulation whereas red crossbars signify inhibition.

The baseline model contains 89 parameters and 40 species. Briefly, each receptor binds to its ligand and forms a receptor-ligand complex that can then become phosphorylated by basally active Src family kinases (SFK). Then, the ligand-bound phosphorylated receptor (pC in **Figure 1**) serves as the catalyst for converting SFK from a basally active state to a fully active state (pSFK)^41^. Next, pSFK mediates the phosphorylation of PLCγ, Vav, SLP76, Akt and Erk, in addition to the phosphatases SHP and SHIP^4,42–46^. The phosphatases SHP and SHIP provide negative feedback to the stimulatory network by dephosphorylating the phosphorylated signaling species, including pC and pSFK. The initial concentrations of the model were taken from the literature^47^, and the kinetic parameters regulating the rate of phosphorylation and dephosphorylation reactions are taken from our published model^19^. In particular, the initial concentrations are assumed to be steady state values since the primary NK cells were pre-incubated in cell culture media which excluded stimulatory ligands for CD16 or NKG2D prior to measurement via mass spectroscopy^47^.

In addition to the model given by Makaryan and Finley^19^, we included a degranulation parameter for each pathway (CD16 and NKG2D) to account for actin remodeling, trafficking, docking and exocytosis of the cytolytic molecules^4,23,42,43,48^. Given pVav’s correlation with NK cell cytotoxicity^41^ and pPLCγ’s role in the release of intracellular calcium ions, which are needed for exocytosis^4,42–44,49,50^, we used pVav and pPLCγ as catalysts for GZMB and PRF1 secretion. Also, data from Srpan, et. al.^27^ demonstrated cross-talk between CD16 and NKG2D; specifically, stimulation of CD16 induced a slight increase in the amount of NKG2D, whereas stimulation of NKG2D slightly decreased the concentration of CD16. We included reactions to account for the crosstalk, based on either mass action kinetics or a nonlinear Hill equation. We also considered alternate models that included synthesis and decay reactions for the inactive signaling species since the data in Srpan, et. al.^27^ stimulated NK cells over a long timescale. Excitingly, all candidate models demonstrated a good agreement with experimental observations. We compared each model using the Akaike information criterion (AIC) which measures the quality of models given a dataset. We found the simpler model, which (1) had linear equations for the crosstalk reactions and (2) excluded synthesis and decay reactions for the inactive species, to be the preferred model (see Table 1). Specifically, we calculated the relative likelihood of the models: 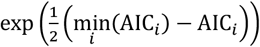, where *i* is the index number of the model. This value represents the how likely or probable the model candidates are to the one with the lowest AIC score.

**Table 1.** AIC scores for model selection

| Model | Crosstalk | Synthesis/Decay Reaction | AIC score | Relative to Model 1 |
| --- | --- | --- | --- | --- |
| 1 | Linear: $kpC(t)$ | N/A | 179.9 | 1 |
| 2 | Nonlinear: $k \frac{pC(t)}{K + pC(t)}$ | N/A | 187.7 | 0.020 times as probable |
| 3 | Linear: $kpC(t)$ | $\emptyset \rightarrow X \rightarrow \emptyset$, where $X$ is inactive species | 188.2 | 0.015 times as probable |

### Data collection and processing

The mathematical model was trained using experimental data from Srpan, et. al.^27^, who used (1) primary NK cells in their experimental studies, (2) Rituximab as the ligand for stimulating CD16 and (3) MICA for the stimulation of NKG2D. The freely available online software WebPlotDigitizer (https://automeris.io/WebPlotDigitizer) was used to extract mean and standard deviation data from plots shown in Srpan, et. al.^27^. In that study, the researchers stimulated primary NK cells in a 96-well plate using immobilized Rituximab and MICA for activation of CD16 and NKG2D, respectively. In addition, each well was coated with anti-PRF1 monoclonal antibodies to measure and visualize the concentration of secreted PRF1 via confocal microscopy. The researchers subsequently quantified the optical density of microscopy images using ImageJ^51^ as a measure of secreted PRF1.

We extracted a total of 50 data points from the published study, from which 34 were used for model training and the remaining 16 were used for model validation. The experimental design in Srpan, et. al.^27^ is as follows: NK cells were stimulated under one pathway for 60 minutes, then washed for 15 minutes prior to the next round or iteration of receptor stimulation. The cells were stimulated for two rounds under one pathway and then either stimulated via the same pathway or the other pathway for the third round. We designated the data from the first two rounds of receptor stimulation for model training, and if the same pathway was stimulated again for the third round, then this data would also be assigned to the training set; otherwise, it would be assigned for model validation. For data where NK cells were stimulated for 120 minutes in a single well (i.e., under one pathway), we used the first half for model training and the second half for model validation. Lastly, since the model predictions are units of concentration (*μ*M) and the experimental data are signal intensity, we normalized both the model predictions and the data prior to training by calculating the percent change (%Δ) from the same reference time point, as done in our previous work^19^:

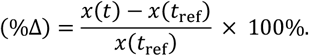

### Parameter estimation

The model parameters were estimated using a Bayesian framework; namely, the Metropolis-Hastings (MH) algorithm^52–55^, as we did previously^19^. We provide an extensive description of this approach in the published work. Briefly, the goal of this approach is to sample from the posterior probability distribution of the parameters given the data (*p*(*θ*|*y*)). Bayes’ theorem provides a relationship between the posterior and prior probability densities via

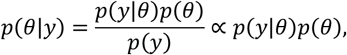

where *p*(*θ*) is our prior knowledge of the parameters, *p*(*y*|*θ*) is the data likelihood function and *p*(*y*) is the probability of the data (which is constant here since the data is given). The seven estimated model parameters are related to secretion and are briefly described in Table 2 below. All other parameters are set to their estimated values from our previous model^19^.

**Table 2.**
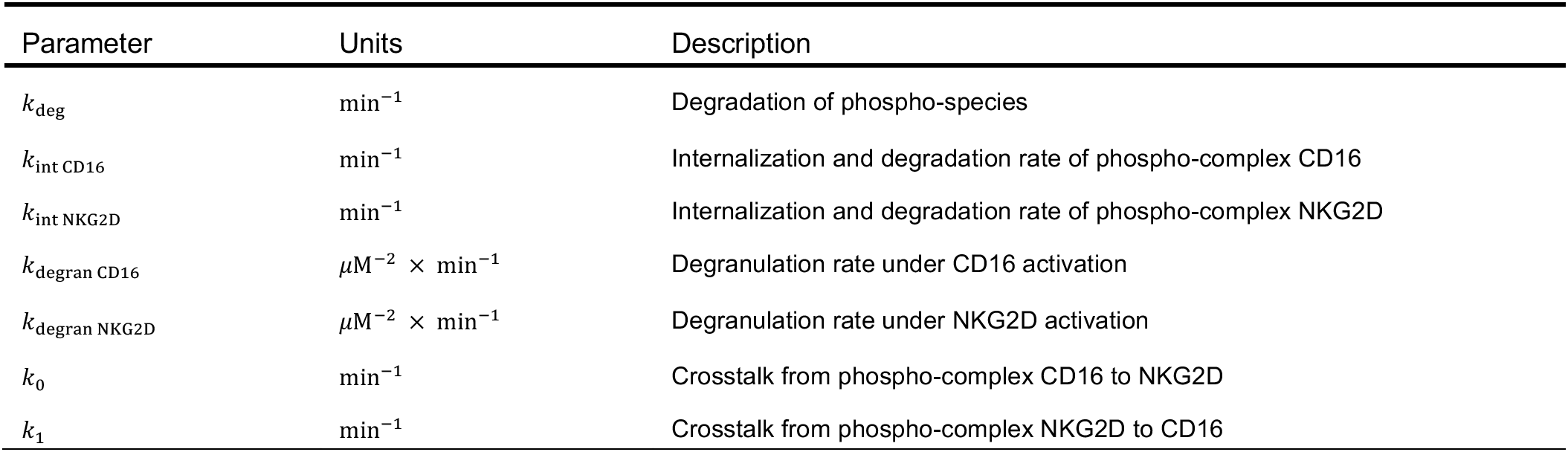
List of estimated parameters

| Parameter | Units | Description |
| --- | --- | --- |
| $k_{\text{deg}}$ | $\text{min}^{-1}$ | Degradation of phospho-species |
| $k_{\text{int CD16}}$ | $\text{min}^{-1}$ | Internalization and degradation rate of phospho-complex CD16 |
| $k_{\text{int NKG2D}}$ | $\text{min}^{-1}$ | Internalization and degradation rate of phospho-complex NKG2D |
| $k_{\text{degran CD16}}$ | $\mu\text{M}^{-2} \times \text{min}^{-1}$ | Degranulation rate under CD16 activation |
| $k_{\text{degran NKG2D}}$ | $\mu\text{M}^{-2} \times \text{min}^{-1}$ | Degranulation rate under NKG2D activation |
| $k_0$ | $\text{min}^{-1}$ | Crosstalk from phospho-complex CD16 to NKG2D |
| $k_1$ | $\text{min}^{-1}$ | Crosstalk from phospho-complex NKG2D to CD16 |

We used a continuous uniform prior distribution on the parameters. Specifically,

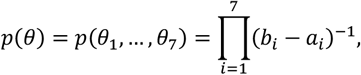

where each *θ*_*i*_ is an element of the interval [*a*_*i*_, *b*_*i*_]. Indeed, the prior distribution independent of *θ*.

The data likelihood function represents a measure of the error between the data and the model. We assume the error between each data point and model prediction 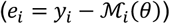 are independent and normally distributed with zero mean and identical variance *σ*^2^:

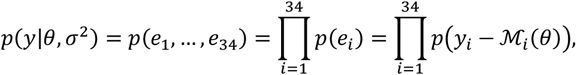

where each 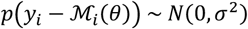 and 34 represents the number of training data. Moreover, we marginalize the noise (*σ*^2^) from *p*(*y*|*θ*, *σ*^2^) by assuming an inverse gamma measure over *σ*^2^ (specifically, *p*(*σ*^2^) ~ Γ^−1^(*α*, *β*)) and integrating *p*(*y*|*θ*, *σ*^2^) with respect to *σ*^2^ to obtain

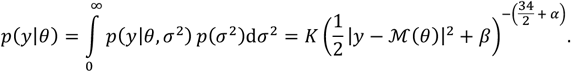

Here, *K* is a constant that is independent of both *y* and *θ*. Note that *p*(*y*|*θ*) achieves its maximum when 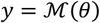, which implies that maximizing the posterior density *p*(*θ*|*y*) is proportional to minimizing the sum of squared residuals.

However, we cannot solve for *p*(*θ*|*y*) analytically due to the nonlinearities in 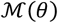. To that end, we employ the MH algorithm to sample from the posterior distribution. Before doing so, we specify key components needed to implement the MH algorithm^53,54^. We use the lognormal distribution as the proposal distribution which proposes a new parameter vector given the current estimate:

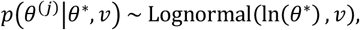

where *θ*^∗^ is the current estimate, *θ*^(*j*)^ is the proposed parameter vector, *j* is the number of iterations of the MH algorithm and *ν* = 0.1 is the scale parameter of the distribution. Next, we compute the acceptance ratio (AR) at each iteration *j*:

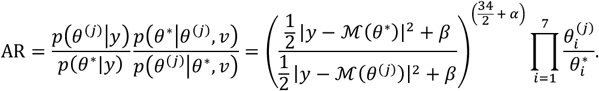

Since *α*, *β* > 0 is needed to satisfy the inverse gamma distribution, we set both equal to one. The last term on the right is the ratio of the transition kernels, which measures the asymmetric probability of transitioning between *θ*^∗^ and *θ*^(*j*)^. Given the above proposal distribution and AR, we simulated the MH algorithm for 10,000 iterations. In our estimation, the marginal posterior distribution of each parameter converges to a stationary distribution anywhere between the 2,000^th^ – 5,000^th^ iteration. Given that the MH algorithm is a stochastic optimization method, we simulated the MH algorithm 200 times using independent, random initial guesses for *θ*^∗^. For simulations, we used the final 1,000 iterations of the MH algorithm which we take to be samples from the posterior distribution. The model is provided in Supplementary File S1 and the list of model species, reactions and parameters are provided in Supplementary File S2.

### Information-theoretic sensitivity analysis

We employed an entropy-based sensitivity analysis for informing which parameters (model inputs) share a significant degree of mutual information with the amount of secreted GZMB and PRF1 (model outputs). We follow the methods described previously by Lüdtke, et. al.^56^. Entropy, in the sense of Shannon, is described as the average information content of a random variable. That is, for any random variable *Y*, the (Shannon) entropy of *Y* is given by

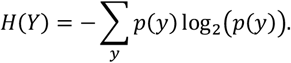

The conditional entropy of *Y* given *X* = *x* (*H*(*Y*|*X* = *x*)) is defined analogously by using the conditional probability *p*(*y*|*X* = *x*). Moreover, the quantity *H*(*Y*|*X*) measures the average amount of information remaining *Y* given that we observed another random variable *X*. Specifically,

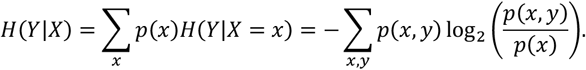

Note that *H*(*Y*) ≥ *H*(*Y*|*X*) as any random variable *X* can only explain away some of the information in *Y*, if any at all. If *H*(*Y*|*X*) = *H*(*Y*), then *Y* is independent of *X*. Alternatively, if *H*(*Y*|*X*) = 0, then knowing *X* completely determines *Y*. As defined in Lüdtke, et. al.^56^, conditional entropies of the form *H*(*Y*|{*X*_1_, …, *X*_*n*_}\*X*_*i*_) measure the total effect a particular input *X*_*i*_ exerts on the output *Y*. Furthermore, the quantity *H*(*Y*|{*X*_1_, …, *X*_*n*_}) determines the amount of information remaining in *Y* once we observed all of the inputs. This can be thought of as the *residual information* that persists in *Y* for which the inputs *X*_1_, …, *X*_*n*_ cannot account for. Thus, the total order sensitivity index for each input *X*_*i*_ is defined by

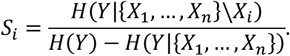

Indeed, inputs with a higher sensitivity index suggest the output is sensitive to variations in the input. In our case, *X*_*i*_ represents the kinetic parameters in our model whereas *Y*_1_ and *Y*_2_ represent the amount of secreted GZMB and PRF1, respectively, after 60 minutes of receptor stimulation to mimic the experimental conditions from Srpan, et. al^27^. We drew 250 random samples for each *X*_*i*_ (independently) using a uniform distribution on the interval 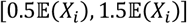, where 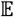 is the expectation operator and the distribution of each *X*_*i*_ is given by parameter estimation (see above) or from Makaryan and Finley^19^. Next, we simulated the model using these 250 samples to generate a distribution for each of the outputs *Y*_1_ and *Y*_2_ to then compute the total order sensitivity indices.

### synNotch signaling and RNA expression model

The synthetic biology field has empowered biologists with tools for engineering novel cellular responses. In general, such molecular programming techniques provide cells with additional capabilities; for instance, Smole, et. al.^57^ engineered novel genetic circuits in mammalian cells to respond to inflammatory signals (i.e., IL-1*β*) by producing anti-inflammatory proteins. In addition, the use of synthetic biology methods with clinical applications is the subject of recent reviews^58–60^. The synthetic Notch (synNotch) signaling pathway^61^, in particular, can be constructed via genetic modifications to trigger a specific cell response when a specific stimulus (e.g., chemical, thermal) is present in the micro-environment. Briefly, the synNotch system includes three components: (1) the synNotch receptor, (2) a transcription factor and (3) a plasmid vector. The extracellular domain of the synNotch receptor can be a single-chain variable fragment (scFv) designed to bind to a specific antigen, similar to chimeric antigen receptors (CARs) that target tumor-specific antigens^16,62,63^. The synNotch receptor and the transcription factor are linked together using peptide sequences that can be cleaved by constitutively expressed membrane proteases once the receptor binds to a specific ligand^61^. Then, the transcription factor becomes unchained from the cell membrane and subsequently free to bind to its promoter site and initiate gene expression.

It follows that if the synNotch receptor’s extracellular domain is engineered to bind to the same ligands as CD16 and NKG2D, then this synthetic pathway will be complementary to the endogenous pathway; that is, the endogenous pathway will mediate the secretion of GZMB and PRF1 while the synthetic pathway will both increase the production of GZMB and PRF1 in addition to enhancing their secretion. To determine if this approach is indeed beneficial, we constructed a mathematical model of the synNotch receptor, based on mass action kinetics, to simulate and predict if such a system necessarily leads to the continuous secretion of GZMB and PRF1 over multiple rounds of stimulation.

We implemented an *in silico* synNotch system to inhibit protein activity. For inhibiting protein activity, the use of long non-coding RNAs (lncRNAs) present new avenues for impeding protein-to-protein interactions in signaling pathways^64–67^. Recently, lncRNA pulldown assays^67–69^, when coupled with mass spectroscopy, have been utilized for identifying novel RNA molecules that can bind to and sequester proteins from signaling. An advantage of protein-sequestering lncRNAs is that, once discovered, they can be sequenced and reverse-transcribed via polymerase chain reaction (PCR) to manufacture its complementary DNA strand (cDNA)^70^. The cDNA can be subsequently integrated into a plasmid vector for expression under the control of the synNotch receptor. In particular, we simulate inhibition of phosphatase activity.

We also apply the synNotch system to promote production of cytolytic molecules. By incorporating a multi-cistronic plasmid downstream of the synNotch receptor, which enables the expression of two or more genes, the production of GZMB and PRF1 can be induced once the NK cell binds to a specific target ligand. This added specificity allows the researcher to designate when protein production to occur. Given that protein expression is a costly function, this strategy limits cytolytic molecule production to situations where an NK cell interacts with a target cell. Thus, the NK cell allocates its resources for cytolytic molecule production only in cases where a target cell is nearby. In contrast to constitutive over-expression techniques, this method allows the NK cell to preserve its energy for other functions.

To simplify our analysis, we assume (1) a lncRNA binds to, and sequesters, both phosphorylated and unphosphorylated phosphatase at a rate of 1 (*μ*M × min)^−1^ and (2) there are two plasmids (at fixed amounts) controlling the expression of lncRNA and the cytolytic molecules separately and that both are under the control of the same transcription factor. For a detailed description of the reactions and parameters that characterize the synNotch signaling system see Table 3.

**Table 3.** synNotch signaling model

| No. | Equation | Reaction Velocities | Parameters | Description |
| --- | --- | --- | --- | --- |
| 1 | $R + L \rightleftharpoons C$ | $k_{\text{on}}R(t)L(t), k_{\text{off}}C(t)$ | $k_{\text{on}}, k_{\text{off}}$ = same as endogenous receptor | Binding kinetics |
| 2 | $C \rightarrow C_* + TF$ | $k_{\text{cleave}}C(t)$ | $k_{\text{cleave}} = 1 \text{ min}^{-1}$ (assumed) | Detachment of TF |
| 3 | $C_* \rightarrow L$ | $k_{\text{int}}C_*(t)$ | $k_{\text{int}}$ = same as endogenous receptor | Ligand recycling |
| 4 | $TF \rightarrow \emptyset$ | $k_{\text{deg}}TF(t)$ | $k_{\text{deg}}$ = same as phospho-protein decay | Degradation of TF |
| 5 | $\emptyset \rightarrow \text{RNA}$ | $k_{\text{trpt}}u \frac{TF(t)}{K_M + TF(t)}$ | $k_{\text{trpt}} = 600 \text{ nt} \times \text{min}^{-1}$ (calculated<sup>71</sup>)<br>$K_M = 1 \text{ min}^{-1}$ (assumed) | RNA production |
| 6 | $\text{RNA} \rightarrow \text{RNA} + \text{protein}$ | $k_{\text{trsl}}\text{RNA}(t)$ | $k_{\text{trsl}} = 600 \text{ aa} \times \text{min}^{-1}$ (calculated<sup>71</sup>) | Protein production |
| 7 | $\text{RNA} \rightarrow \emptyset$ | $k_{\text{deg RNA}}\text{RNA}(t)$ | $k_{\text{deg RNA}} = 0.0462 \text{ min}^{-1}$ (calculated<sup>72,73</sup>) | RNA degradation |

In equation (1), the ligand can be either Rituximab or MICA, to signal complementarily with the CD16 or NKG2D pathway, respectively. For equation (5), the production of RNA is non-linear with respect to the transcription factor (TF) and follows a Hill function with plasmid affinity *K*_M_ and Hill coefficient equal to one. We assume the plasmid concentration (u) in equation (5) is at steady state prior to receptor stimulation. Additionally, we performed a sensitivity analysis to determine the robustness of the model predictions with respect to our assumptions on the kinetic rates.

### Optimization of synNotch system

We apply optimal control theory to determine how a synthetic pathway, the synNotch pathway, can be used to promote maximal secretion of cytolytic molecules. Such an optimization is necessary because, while the synNotch pathway may lead to an increase in the production of cytolytic molecules and the inhibition of protein activity, it certainly imposes a burden on the cell to express this system. Moreover, given that the synNotch and endogenous receptors will be competing for the same ligand, it is not immediately clear how much synNotch receptor is optimal for a given frequency of NK cell stimulation. Therefore, we considered optimizing the synthetic pathway to maximally secrete GZMB and PRF1 while using the absolute minimal amount of exogenous material. In this context, the plasmids and synNotch receptor can be considered as the controllers for the secretion of cytolytic molecules. Thus, our objective is to find the optimal values of the plasmids and synNotch receptor (controllers) such that we maximally induce secretion (performance) at minimal cost to the cell (effort). Indeed, this is an optimization problem, which we can solve using conventional methods^74–77^; specifically, we minimized the following stochastic objective function, given the model parameters *θ*:

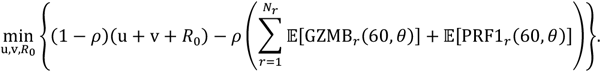

Indeed, we minimize the objective function using the sample average approach (SAA)^78–81^. Here, u, v and *R*_0_ represent the amount of the lncRNA-coding plasmid, cytolytic molecule-coding plasmid and the initial value of the synNotch receptor, respectively. The minimization is subject to the constraints:

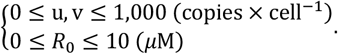

We set the upper bound of the plasmid concentrations based on what is defined as a high copy number^57,60,61^. In addition, the upper bound on the initial value of synNotch receptor is based on a study^82^ where Chinese hamster ovary (CHO) cells were genetically modified to maximally produce human IgG; the values ranged from 0.3 – 20 *μ*M, from which we chose 10 *μ*M. While we acknowledge the dissimilarities between the CHO and NK cells, and the synNotch receptor and human IgG, it is nevertheless a strict upper bound that can constrain our estimation.

The second term on the right represents the cumulative secretion of GZMB and PRF1 after 60 minutes of receptor stimulation, where *r* = 1, …,*N*_*r*_ is the number of rounds of receptor stimulation and the expectation is taken over the parameters *θ*. Since GZMB and PRF1 are non-negative, the second term is in fact a maximization problem given that argmin −*f* = argmax *f* for all non-negative *f*. The constant *ρ* ∈ (0,1) is a weight parameter that specifies how much emphasis is placed on minimizing the first term (effort) versus maximizing the second term (performance); in this study, we place equal weight on both (i.e., 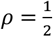).

To solve this optimization problem, we used the mesh-adaptive directed search (MADS) algorithm *patternsearch* in MATLAB. This is a gradient-free method that attempts to locate a minimizer of the objective function by evaluating many trials points nearby the initial guess at each iteration. If some trial point near the initial guess induces a lower function evaluation, then the iteration terminates and starts again by implicitly creating a new mesh around this new incumbent point. If the algorithm cannot find a feasible point that minimizes the objective function, the mesh around the current incumbent point becomes finer and finer until a predefined threshold is reached. For our purposes, we set the mesh tolerance parameter to 10^−6^ at which the algorithm terminates.

## 3 RESULTS

### 3.1 NK cell degranulation model can reproduce experimental observations

We generated a mathematical model of NK cell degranulation by incorporating the dynamics of GZMB and PRF1 into our previous model of NK cell signaling^19^. In total, the model consists of downstream signaling reactions from the receptors CD16 and NKG2D that ultimately mediate the secretion of the cytolytic molecules GZMB and PRF1. Specifically, the model consists of a system of nonlinear ODEs that predict the concentration of the receptors, signaling intermediates and the cytolytic molecules; explicit equations, initial conditions and parameters of the model are provided in Supplementary File S2. When the CD16 and NKG2D receptors are stimulated, they activate the cell via a cascade of reactions (**Figure 1**): activation of the Src family kinases (pSFK), facilitated by the ligand-bound phosphorylated receptors (pC), mediates the activation of the Akt, SLP76-Vav-Erk, and PLCγ pathways. In particular, we assume pVav and pPLCγ mediate the secretion of GZMB and PRF1 as they have been correlated with NK cell cytotoxicity^41^ and a discharge of intracellular calcium ions^9,10,43,44,49^ which mediate exocytosis, respectively. Although Akt and Erk are necessary for cell survival and proliferation^9,23,42,44^, respectively, their contribution to NK cell secretion of GZMB and PRF1 remains elusive. As it stands, given the available data, we consider pVav and pPLCγ as the mediators for NK cell secretion.

The model was calibrated to data from Srpan, et. al.^27^ using a Bayesian perspective to parameter estimation^53,54^; namely, we implemented the Metropolis-Hastings (MH) algorithm (see methods) to sample from the posterior distribution of the parameters conditional on the data. In brief, seven parameters (see Table 1) were estimated 200 times using randomized initial guesses. We also tested the model predictions using a separate validation dataset. The combined error for each run can be found in **Figure S1**. Since the marginal posterior distribution of each parameter for the best twenty runs were almost identical, we chose to simulate the model using the results from the best run (Run 1 in **Figure S1**). The trace plots for each parameter in the best run are shown in **Figure S2**, where the value of each parameter is plotted as a function of the iteration of the MH algorithm. This diagnostic of the parameter estimation shows that each parameter converges to a stationary distribution, albeit at different iterations. We simulated the model 1,000 times using the final 1,000 iterations of each parameter from the best run (**Figure S3**) and compared the results to both the training and validation data.

Excitingly, the model simulations are in good agreement with both the training and validation data (**Figure 2**). The model can reproduce the time evolution of secreted PRF1 mediated by stimulation of NKG2D (**Figure 2A**) and CD16 (**Figure 2B**). In addition, the model can replicate sequential stimulation data where NK cells are stimulated via one pathway for two consecutive rounds, each for 60 minutes, and then stimulated via the other pathway for the third round of stimulation. Under these conditions, the simulated concentration of secreted PRF1 (**Figures 2C – 2D**), the receptors (**Figures 2E – 2H**) and intracellular PRF1 (**Figures 2I – 2J**) all agree with the experimental data. To further validate the model predictions, we simulated the concentration of secreted PRF1 when both Rituximab and MICA were decreased to 1 *μ*g/mL (**Figure 2K**), and these model simulations agree with experimental measurements as well. Overall, we generated an experimentally validated, mechanistic model of NK cell degranulation, which can recreate results from Srpan, et. al.^27^ under a range of different experimental conditions. We next apply the model to determine which features, when perturbed, robustly maximize the amount of secreted PRF1 and GZMB.

**Figure 2.**
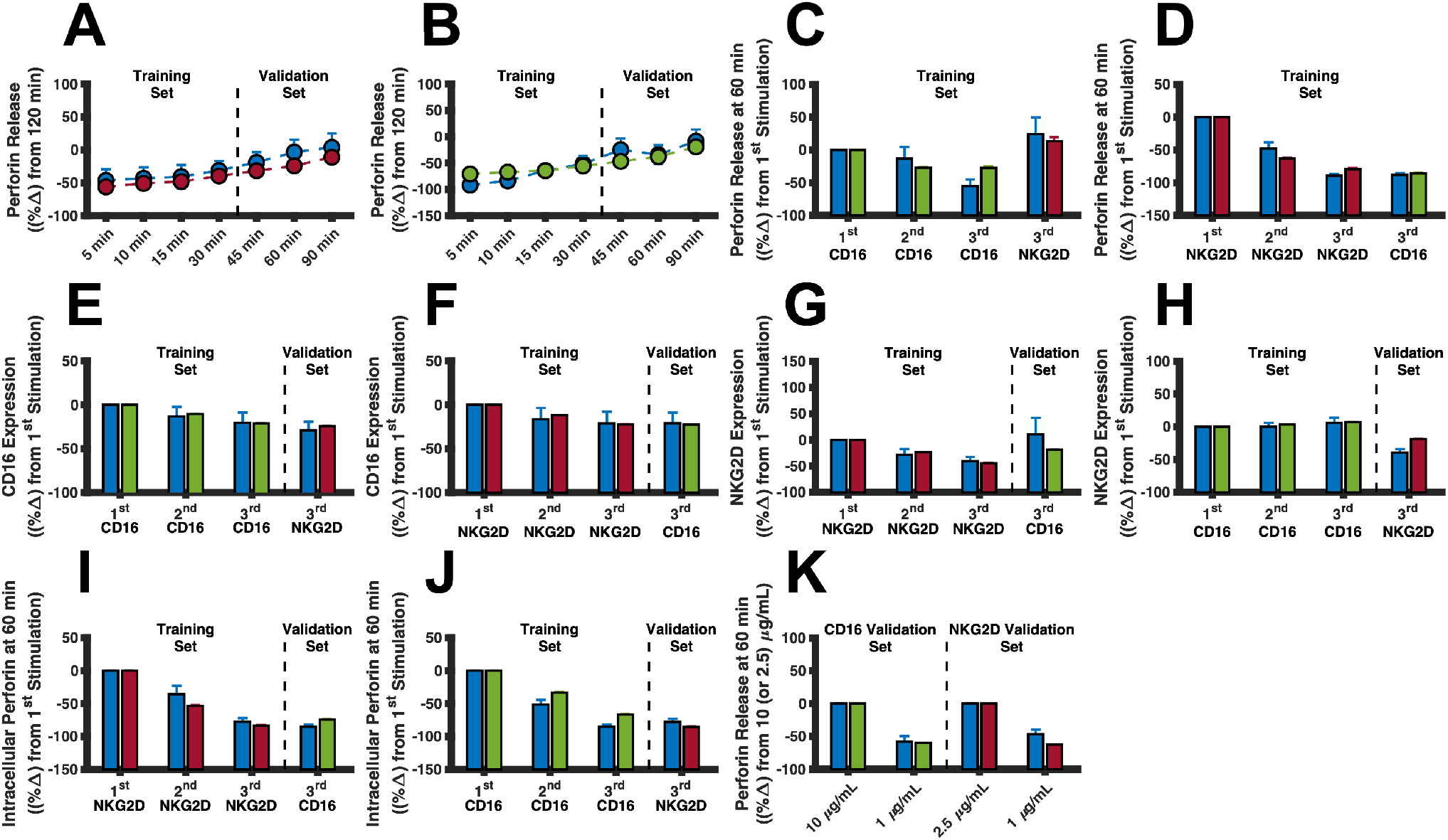
Model training and validation. The model was trained to, and tested against, *in vitro* NK cell stimulation data from Srpan, et. al. (2018). In all panels, blue markers and bars represent mean experimental data, while red and green signify the mean model predictions for the NKG2D and CD16 pathways, respectively. Error bars indicate standard deviations. Except for panel (**K**), the ligand concentration used to stimulate NKG2D and CD16 are 2.5 *μ*g/mL of MICA and 10 *μ*g/mL of Rituximab, respectively. (**A** and **B**) Normalized time series data of PRF1 secretion using (**A**) 2.5 *μ*g/mL of MICA or (**B**) 10 *μ*g/mL of Rituximab. (**C** and **D**) Normalized PRF1 secretion per round of stimulation by stimulating the CD16 or NKG2D pathway and then stimulating the NKG2D pathway or CD16 pathway. (**E** and **F**) Normalized concentration of CD16 per round of stimulation by stimulating the CD16 or NKG2D pathway and then stimulating the NKG2D or CD16 pathway. (**G** and **H**) Normalized concentration of NKG2D per round of stimulation by stimulating the CD16 or NKG2D pathway and then stimulating the NKG2D or CD16 pathway. (**I** and **J**) Normalized concentration of intracellular PRF1 by stimulating the NKG2D or CD16 pathway and then stimulating the CD16 or NKG2D pathway. (**K**) Normalized concentration of PRF1 after 60 minutes of stimulation using different concentrations of ligand.

### 3.2 Inhibition of pSHP maximizes PRF1 and GZMB secretion *in silico*

In order to understand which model parameters (inputs) leads to an increase in the predicted secretion of GZMB and PRF1 (outputs), we performed an entropy-based sensitivity analysis on the model (see methods). This technique, as shown by Lüdtke, et. al.^56^, measures the degree of mutual information shared between the model inputs and outputs. The greater the amount of shared mutual information between an input and output, the more sensitive the output is to that input. This metric is defined as the sensitivity index of the parameter (see methods). Briefly, we varied the model parameters 50% above and below their mean value, then drew 250 uniformly distributed samples to simulate the model and generate distributions for the amount of GZMB and PRF1 after 60 minutes of receptor stimulation. With these probability distributions for the model inputs and outputs, we are able to approximate the conditional entropies needed to compute the sensitivity indices.

Interestingly, only a select few parameters were shown to have a large sensitivity index in regard to GZMB and PRF1 secretion (**Figure S4**). The fifteen most influential parameters are found in the pSFK-pSHP-pVav-pPLCγ subgraph, where each parameter can explain more than 35% of the information in GZMB secretion promoted by the CD16 receptor (**Figure 3A**; **Figure S4A**). This also holds true for NKG2D-mediated GZMB secretion (**Figure 3B**; **Figure S4C**) as well as for PRF1 secretion promoted by either receptor (**Figures S4B, D**). These results suggest that this subnetwork (**Figure 3C**) strongly influences the amount of GZMB and PRF1 secretion. In fact, the total effect of the most influential parameter, describing the catalytic rate of pSHP activation, shares over 70% of the information observed in GZMB and PRF1 secretion. Given the causal structure of the model, we can infer that blocking the pSFK → pSHP reaction can yield more GZMB and PRF1 secretion since that would prevent the phosphatase from inhibiting signal transduction. Similar causal inferences can be made for other parameters in the subgraph in **Figure 3C**. Thus, we intervened on these parameters (individually) by modifying their values and simulating the model to measure the percent change in GZMB and PRF1 secretion from baseline compared to the modified case.

**Figure 3.**
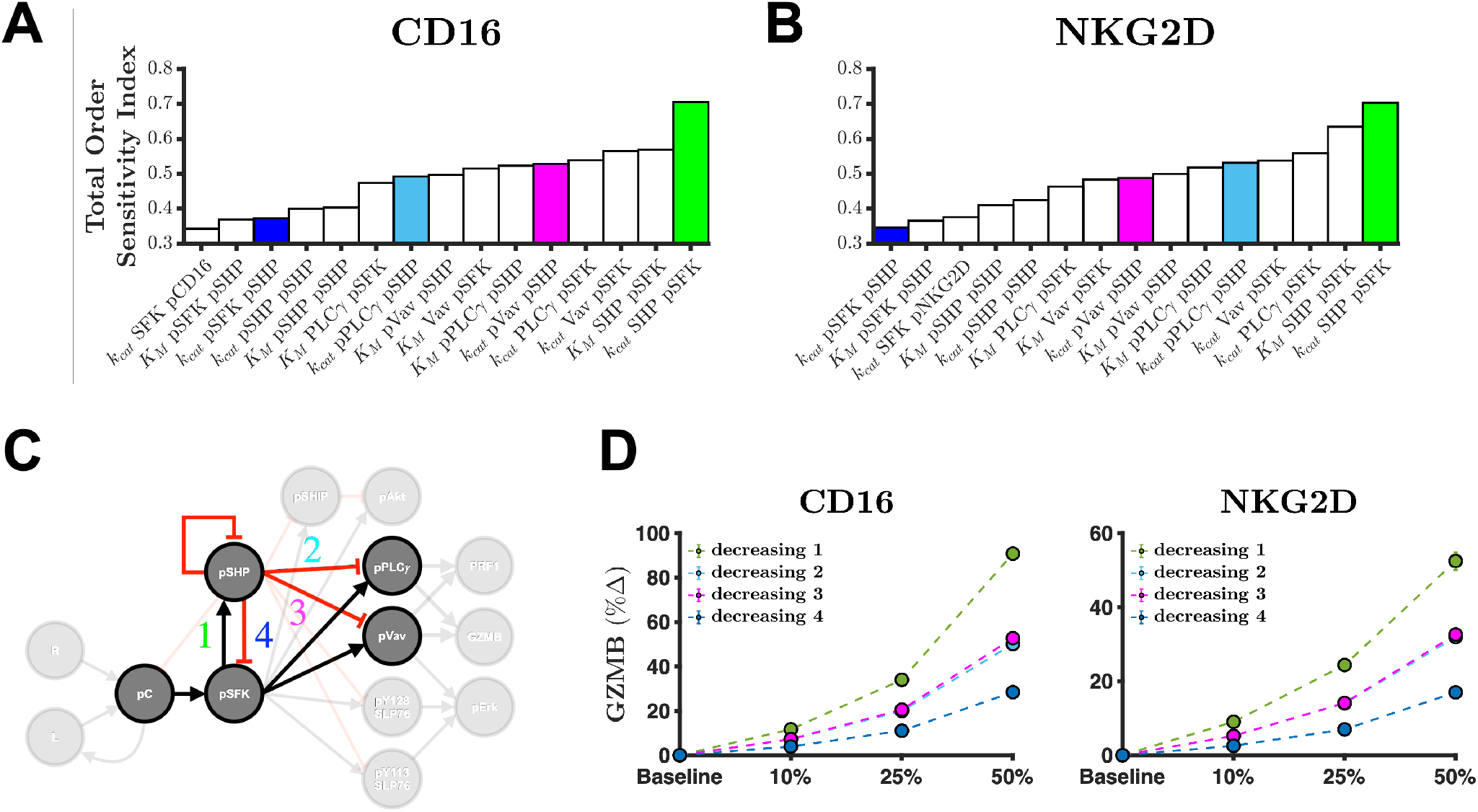
Sensitivity analysis shows inhibition of pSHP activation is most influential for GZMB and PRF1 secretion. Total order sensitivity indices of the top fifteen influential parameters for the amount of GZMB secretion after 60 minutes of stimulation of the (**A**) CD16 pathway or (**B**) NKG2D pathway. (**C**) Almost all of the influential parameters from the sensitivity analysis can be found in the pSFK-pSHP-pPLCγ-pVav sub-network of the full model. Predicted percent change in GZMB secretion when decreasing the catalytic rate constants (*k*_cat_) parameters involved in the sub-network depicted in (**C**) after 60 minutes of stimulation of (**D**) CD16 and (**E**) NKG2D. In panels (**D** – **E**): green, decreasing pSHP activation; blue, decreasing of pSFK deactivation; magenta, decreasing pVav deactivation; cyan, decreasing pPLCγ deactivation; markers, mean model prediction; error bars, one standard deviation.

Since there are four variables of interest in this influential sub-network (pSFK, pSHP, pVav, pPLCγ), each with a rate of activation and deactivation, we intervened on eight different parameters to enhance or inhibit the phosphorylated species. Specifically, the catalytic rate constants were varied from baseline. Then, the model was subsequently simulated to quantify their impact on the percent change in GZMB and PRF1 secretion after 60 minutes of receptor stimulation (**Figure 3D and Figure S5**). The simulated percent increase in GZMB secretion is greatest when the pSFK → pSHP reaction is inhibited (**Figure 3C**; decreasing 1), when either CD16 or NKG2D (**Figure 3D**) is stimulated. The model predicts that reducing the catalytic rate constant for this pathway has a bigger effect when CD16 is stimulated, compared to NKG2D stimulation (**Figure 3D**). The remaining interventions were not as influential, however (**Figures S5A and S5D**). Furthermore, this conclusion is equally true for PRF1 secretion (**Figures S5B, S5C, S5E and S5F**). These results suggest that disrupting the incoherent feed-forward loop (IFFL) pSFK → pSHP ⊣ pX, where X is either Vav or PLCγ, leads to more GZMB and PRF1 secretion. In summary, we interrogated the model and found that the amount of cytolytic molecule secretion is strongly affected by the pSFK → pSHP edge in the subgraph depicted in **Figure 3C**.

### 3.3 The optimal synNotch system depends on the number of rounds of stimulation

The results above indicate that a strategy to increase GZMB or PRF1 secretion is to inhibit pSHP activation. While this does lead to more GZMB and PRF1 secretion *in silico*, it almost completely depletes the intracellular pool of cytolytic molecules (**Figure S6**). Depleting intracellular pools of GZMB and PRF1 makes the NK cell less likely to secrete these cytolytic molecules upon subsequent stimulation since the time-scale over which the pool is replenished (i.e., protein production) is longer than the timescale over which the cell would be stimulated^40,71^. Ultimately, this causes NK cells to become less cytotoxic over time. Therefore, to maximize GZMB and PRF1 secretion over multiple rounds of stimulation, we need to induce the production of these molecules, in addition to reducing SHP activation. Fortunately, the field of synthetic biology^58–60^ provides tools that can be applied to address this issue. Multi-cistronic plasmids, which express two or more genes, can be used to promote expression of both the GZMB and PRF1 genes in NK cells and thereby replenish the intra-cellular pool of the cytolytic molecules. When coupled with the inhibition of pSHP, this strategy may increase both cytolytic molecule secretion and production, enabling NK cells to continuously secrete cytolytic molecules over multiple rounds of stimulation.

Excitingly, the synthetic Notch (synNotch) signaling system^61^ can be applied to simultaneously (1) inhibit pSHP and (2) increase the intracellular pools of the cytolytic molecules when the NK cell senses a danger signal (e.g., MICA). In brief, the synNotch system can be constructed via genetic modifications. Once the cell expresses the synNotch receptor, it is able to promote a specific function^58–60,83^ that can be tailored to a specific stimulus (**Figure 4A**). We apply this framework as a mechanism for controlling the production and secretion of the cytolytic molecules (i.e., the specific functions) when the NK cell binds to CD16 and NKG2D ligands (i.e., the specific stimuli). For inhibiting pSHP, we simulate the expression of long non-coding RNA (lncRNA) targeting SHP, which impedes SHP from binding to its targets, thereby reducing its ability to inhibit signaling^64,65,67^ (see methods). At the same time, we consider increasing the expression of GZMB and PRF1 using the synNotch system. Collectively, this approach can grant NK cells the ability to maximally secrete cytolytic molecules upon repeated receptor stimulation.

**Figure 4.**
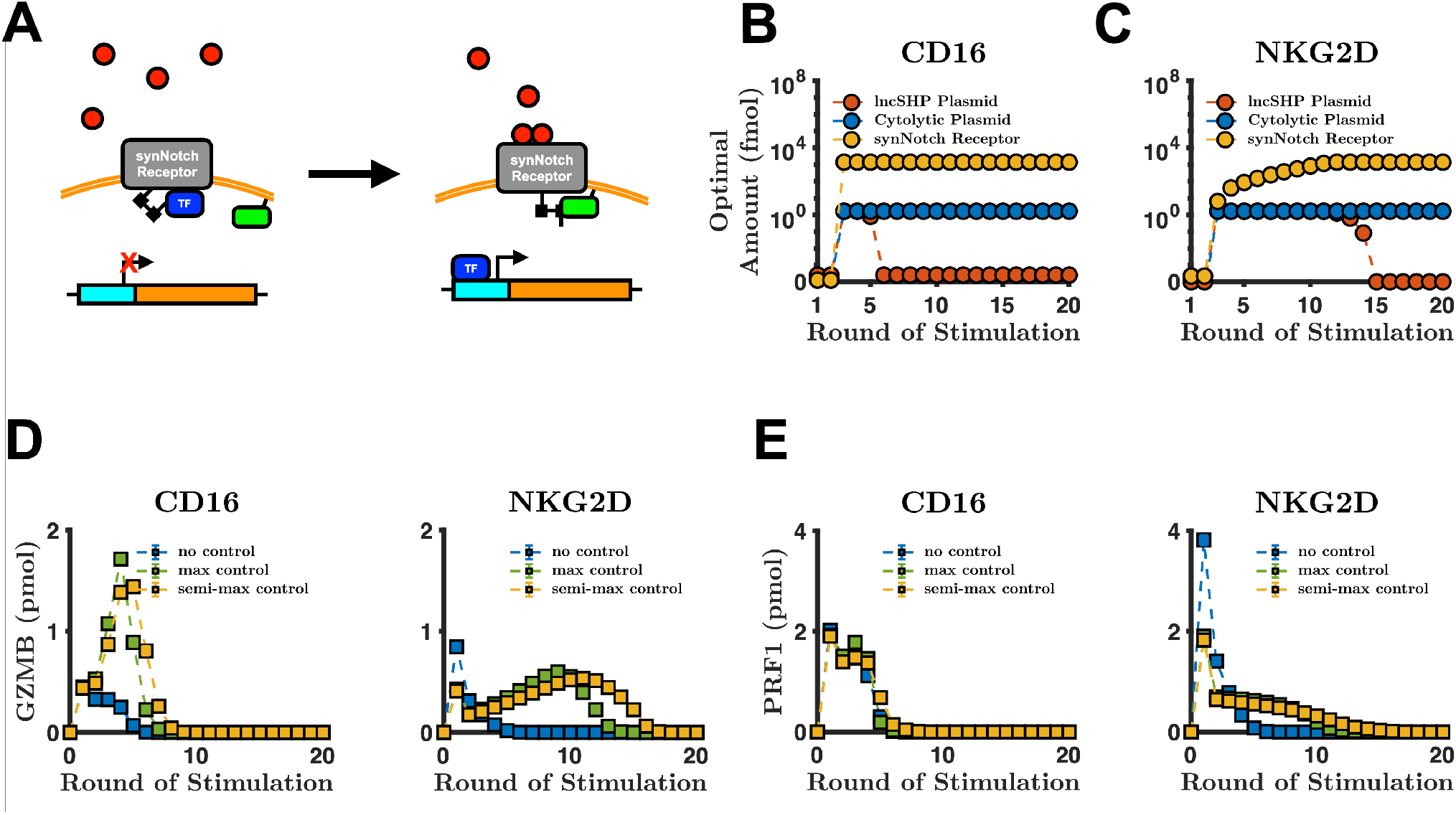
Optimal strategies depend on the number of rounds of stimulation. (**A**) Diagram of synNotch signaling pathway. The ligands (red) can bind to synNotch receptor, which allows the complex to change in conformation to allow constitutively expressed, membrane-bound proteases (green rectangle) to cleave the peptide link between the synNotch receptor and the transcription factor (TF, blue). This allows the TF to bind to its binding site (cyan) on the plasmid which initiates gene (brown) transcription. The objective function is minimized over various rounds of stimulation of (**B**) CD16 and (**C**) NKG2D. For each round of stimulation, the optimal amount of lncSHP plasmid (red markers), cytolytic molecule plasmid (blue markers) and synNotch receptor (orange markers) are shown. (**D – E**) The performance of the NK cell with and without controllers. The secretion of (**D**) GZMB and (**E**) PRF1 via the CD16 or NKG2D pathway. “No control,” optimal values from round 1 for CD16 and NKG2D; “max control,” optimal values from round 3 for CD16 and round 11 for NKG2D; “semi-max control,” optimal values from round 6 for CD16 and round 15 for NKG2D. Markers, mean model prediction; error bars, one standard deviation.

We study the case where the extracellular domain of the synNotch receptor has the same binding kinetics as the endogenous receptor (see Table 3). In this case, the synthetic and endogenous pathways are independent yet complementary to one another: the synthetic path produces the cytolytic molecules while the endogenous pathway secretes them. Although this strategy seems intuitive, the optimal amount of synNotch receptor is difficult to deduce *a priori* given the competing dynamics for ligand. Moreover, it is unclear how much of each plasmid is required to maximally secrete GZMB and PRF1 when considering the number of rounds of stimulation. Since transcription and translation are demanding biological processes^57,60,82^, which are required to express the synNotch receptor in addition to the lncRNA and cytolytic molecules, we consider using the absolute minimal amount of exogenous material needed to achieve maximum secretion. Taken together, we aim to find the optimal levels of lncRNA-coding plasmid targeting SHP (lncSHP), cytolytic molecule-coding plasmid and the synNotch receptor needed to maximize GZMB and PRF1 secretion over many rounds of stimulation while using the absolute minimal amount of synthetic material (see methods).

Interestingly, the optimized synNotch system has different characteristics when paired with CD16 (**Figure 4B**) or NKG2D (**Figure 4C**). When CD16 is stimulated for 1 – 2 rounds only (**Figure 4B**), it is optimal to do nothing. That is, the amount of effort required to express the synNotch system outweighs the gain in performance (i.e., increased secretion). This holds true for NKG2D as well (**Figure 4C**). In contrast, when CD16 is stimulated for 3 – 5 rounds, the optimal amounts for each plasmid and *R*_0_ (initial value of synNotch) are at their maximum values. At this stage, the synNotch system is fully operational. Surprisingly, as we continue to increase the number of rounds of CD16 stimulation, we found that the optimal amount of lncSHP-coding plasmid decreases precipitously by several orders of magnitude, while the optimal amounts of the cytolytic molecule-coding plasmid and *R*_0_ remain at their maximum values. Intriguingly, the model prediction reveals that inhibition of SHP is not optimal in the long-term, as the optimal amount of the lncSHP plasmid is at its minimal value of zero. This is because too much inhibition of SHP leads to an accumulation of phospho-proteins, which increases the velocity of phosphoprotein decay^42,84,85^ (see Table 2, *k*_deg_). Moreover, since there is no synthesis reaction of the inactive signaling species in our model (see Table 1), the degraded phospho-species reduces the total amount of available molecules for the next round of signaling. Thus, for many rounds of stimulation, the phosphatase counterintuitively is needed to maintain the availability of proteins for subsequent signaling.

The optimal synNotch system when coupled with NKG2D has subtle yet important differences. The model predicts that when NKG2D is stimulated for three rounds or more, the optimal amount cytolytic molecule-coding plasmid is at its maximum (**Figure 4C**) – similar to CD16 (**Figure 4B**). Unlike CD16, however, the optimal amount of lncSHP-coding plasmid remains at maximum value from 3 – 14 rounds of stimulation, and the optimal concentration of *R*_0_ does not immediately reach its maximum value. This difference between the optimal amounts of *R*_0_ is due to the initial amounts of the CD16 and NKG2D receptors (38 and 0.3 *μ*M, respectively), compared to the maximum value *R*_0_ can have, 10 *μ*M^82^ (see methods). We interrogated the model and found that the concentrations of CD16 and NKG2D leads to this distinction. The high concentration of the endogenous CD16 receptor compared to the concentration of the synNotch receptor means that CD16 will outcompete the synNotch receptor. This helps clarify why the optimal value of *R*_0_ is the maximal value it can take on. In comparison, for NKG2D stimulation, where the initial amount of the endogenous receptor is less than that of the maximal amount of *R*_0_, a large *R*_0_ would lead to less secretion by directing the input signal more towards cytolytic molecule production. Moreover, since the optimal *R*_0_ is small when considering 10 or fewer rounds of NKG2D stimulation, the indirect impact of SHP inhibition on phospho-protein decay is also small, thus allowing the amount of lncSHP-coding plasmid to remain at maximum for more rounds of NKG2D stimulation. Nevertheless, as the rounds of NKG2D stimulation increases, where the optimal *R*_0_ approaches its maximal value, the optimal amount of lncSHP-coding plasmid reduces by several orders of magnitude – similar to the case for CD16 stimulation. In summary, we not only found that the optimal synNotch system depends on the number of rounds of receptor stimulation but is also different when considering whether the CD16 or NKG2D pathway is being stimulated.

### 3.4 The predicted optimal CD16- and NKG2D-synNotch pairs show qualitative differences in cytolytic molecule secretion

Given the optimized synNotch system, we next simulated the model to predict how the secreted amount of GZMB and PRF1 changes, compared to the baseline case without the synNotch receptor. We simulated the model using the optimal amounts of each plasmid and the initial synNotch receptor from different rounds of stimulation to observe any differences in GZMB and PRF1 secretion. The results of our optimization analysis show three separate clusters of optimal conditions, which depend on the number of rounds of receptor stimulation: (1) no synNotch system, (2) maximal amount of both plasmids and synNotch, and (3) no lncSHP-coding plasmid but maximal amount of cytolytic molecule-coding plasmid and synNotch. For simplicity, we label these clusters as “no control,” “max control” and “semi-max control,” respectively. For CD16, we used the optimal amounts from **Figure 4B** for rounds 1, 3 and 6, respectively, for these three conditions; whereas for NKG2D, we used the optimal amounts from **Figure 4C** for rounds 1, 11 and 15, respectively.

When considering GZMB secretion (**Figures 4D**), the model analysis shows that using the amounts of the synNotch receptor and plasmids optimized for the one round of stimulation (in this case, none at all), more GZMB is released compared to using the amounts of the receptor and plasmids optimized for multiple rounds of stimulation in the first few rounds of stimulation. Specifically, not having the synNotch receptor is best for up to one round of stimulation of the CD16 pathway and up to two rounds of stimulation of the NKG2D pathway. This counterintuitive result that no control is optimal is due to the competition for ligand between the endogenous and the synNotch receptors. For short-term stimulation, it is best for the NK cell to rely on endogenous signaling to maximize GZMB and PRF1 secretion (**Figures 4D, E**). These simulated results help contextualize why it was optimal to not intervene in the first few rounds of stimulation (**Figures 4B, C**).

Interestingly, when the synNotch system is absent, we found that one round of NKG2D stimulation leads to much more secreted PRF1 than CD16 stimulation (3.8 versus 2.0 pmol, respectively), corroborating the findings in Srpan, et. al.^27^. However, for the second round of stimulation, the amount of secreted PRF1 reduces significantly for NKG2D stimulation to 1.4 pmol while CD16 stimulation yields 1.5 pmol of secreted PRF1. For the subsequent round of stimulation, NKG2D yields 0.8 pmol whereas CD16 produces 1.5 pmol. This initial large burst in NKG2D-mediated PRF1 secretion can be explained by the estimated degranulation parameter *k*_degran NKG2D_ (**Figure S3**), which is approximately three times larger than *k*_degran CD16_, and thus the velocity of secretion is faster via the NKG2D path. Taken together, in the absence of a synthetic pathway, NKG2D leads to a large but transient secretion of PRF1 whereas CD16 produces a small but steady secretion of PRF1 after two rounds of stimulation *in silico*.

In contrast, when we consider more rounds of stimulation, it is best to use the synNotch system. Using the “max control” set (rounds 3 and 11 for CD16 and NKG2D stimulation from **Figure 4B – C**, respectively) leads to more GZMB secretion for 2 – 4 rounds of CD16 stimulation and for 3 – 10 rounds of NKG2D stimulation (green squares in **Figure 4D**) when compared to the “no control” set (blue squares in **Figure 4D**). Lastly, the “semi-max control” set (rounds 6 and 15 for stimulation of CD16 and NKG2D from **Figure 4B – C**, respectively) secretes more GZMB after 5 rounds of CD16 stimulation and after 11 rounds of NKG2D stimulation when compared to the “max control” set (compare green to orange squares in **Figures 4D, E**). As a result, the “semi-max control” set is predicted to be the preferred strategy when the number of rounds of stimulation continues to increase. These conclusions hold for PRF1 secretion also (**Figure 4E**). The model again predicts differences between the CD16-synNotch pair and the NKG2D-synNotch pair. When using the “max control” set, the secreted amount of GZMB mediated by the CD16-synNotch pair decreases to zero after 7 rounds of stimulation (**Figure 4D**). The NKG2D-synNotch pair, on the other hand, continues to secrete GZMB up to 14 rounds of stimulation. The model predicts similar differences when considering secreted PRF1, where secretion of PRF1 promoted by stimulation of CD16 goes to zero after 8 rounds of stimulation. In contrast, PRF1 secretion promoted by NKG2D stimulation goes to zero after 16 rounds of stimulation. The differences in concentration between synNotch and the endogenous receptors are mainly responsible for the differences in the responses. Given that *R*_0_ is greater than the initial value of NKG2D (0.3 *μ*M), more of the signal is veered towards cytolytic molecule production due to competition for ligand. This explains NKG2D-synNotch’s sustained response to stimulation. The initial concentration of CD16 (38 *μ*M), in contrast, is greater than *R*_0_, and therefore more of the signal is shifted to cytolytic molecule secretion, resulting in a large but transient response. Taken together, in the presence of synNotch, the qualitative features of CD16-versus NKG2D-mediated secretion of the cytolytic molecules changed compared to the “no control” case: the CD16 path now produces a large but transient response to stimulation while the NKG2D pathway yields a small but sustained response to stimulation.

Interestingly, the optimal synNotch system does not indefinitely increase the amount of secreted cytolytic molecules as we increase the frequency of NK cell stimulation. In fact, when the number of rounds of receptor stimulation becomes large (**Figures 4D, E**), the secreted amount of the cytolytic molecules decreases. We interrogated the model to better understand this prediction. While the intracellular pool of GZMB (**Figure S7A**) and PRF1 (**Figure S7B**) continues to increase as we increase the number of rounds of stimulation, the intracellular amount of pPLCγ (**Figure S7C**) and pVav (**Figure S7D**) eventually decrease to zero. Since the secretion of GZMB and PRF1 is mediated by pPLCγ and pVav in our model, it follows that once the amount of the pPLCγ and pVav becomes negligible, so too does the secretion of GZMB and PRF1. Thus, at higher frequencies of stimulation, the depletion of intracellular signaling species is predicted to be a limiting factor in cytolytic molecule secretion.

To assess the robustness of the results above, we performed a sensitivity analysis on the optimized model that includes the synNotch pathway (see Table 3). Similar to our previous sensitivity analysis, we varied each parameter by 50% above and below its baseline value to create a uniform distribution. Next, we drew 250 samples and simulated the model to determine how sensitive the secretion of GZMB and PRF1 is to the parameters under stimulation of CD16 (**Figure S8**) or NKG2D (**Figure S9**). This sensitivity analysis was performed using the “max control” set (3 and 11 rounds of stimulation for CD16 and NKG2D, respectively) and the “semi-max control” set (6 and 15 rounds of stimulation for CD16 and NKG2D, respectively) optimal results from **Figures 4B – 4C**. For CD16, we found that the total order sensitivity indices of each of the parameters in the synNotch system, for GZMB secretion from the optimal results for the “max control” set (round 3), can at most explain 32% of the information in the output (**Figure S8A**). Moreover, when compared to the parameters in the endogenous pathway (**Figure S8A**; orange versus black), the synNotch parameters are much less relevant, meaning the secretion of GZMB is robust to our assumptions related to the synNotch signaling model. The parameters found to be the most influential are the same as those found in **Figure 3A**, which belong to the endogenous pathway. Similarly, these conclusions hold for the “semi-max control” case of GZMB secretion (**Figure S8C**) as well as for PRF1 secretion (**Figure S8B, D**) and GZMB and PRF1 secretion promoted by the NKG2D pathway (**Figure S9**). To further verify these results, we intervened on the *R*_0_ and *k*_on_ parameters in the synNotch pathway (which are the most influential parameters from the synNotch system) as we presented above (**Figure 3D**) and compared the results to decreasing the activation rate of pSHP (**Figure S10**). We found no significant change in the secretion of GZMB or PRF1. In conclusion, we augmented our baseline model with a synNotch signaling system and found that the secretion of the cytolytic molecules can be enhanced with this added system, and the predicted impact of the synNotch system is not significantly affected by the parameters and assumptions we implemented.

## 4 DISCUSSION

In the present work, we constructed a mathematical model of NK cell secretion of GZMB and PRF1, which replicated experimental observations from Srpan, et. al.^27^. The model was simulated to better understand which subnetworks strongly regulate the secretion of the cytolytic molecules as well as strategies for maximizing their secretion *in silico*. Furthermore, we investigated *in silico* how to enhance secretion of the cytolytic molecules over time. Specifically, we simulated the effects of engineering the NK cell to express the synNotch receptor that simultaneously enables production of GZMB and PRF1 and expression of an inhibitor of the phosphatase that significantly affects GZMB and PRF1 secretion. Our modeling revealed the following: as the number of rounds of stimulation increases, the optimal initial value of the synNotch receptor needed to increase proportionally. This implies that if the NK cell is to be stimulated for multiple rounds, it is optimal to promote the synthetic pathway in order to increase production of GZMB and PRF1. In general, the production of the cytolytic molecules should be induced as maximally as possible. The optimal amount of lncSHP-coding plasmid, however, should switch from its maximum value to the minimum (0 copies per cell) as the number of rounds of stimulation increases, suggesting SHP inhibition is only optimal in the short term. In total, our work presents a theoretical framework that can be used by researchers for engineering NK cells with applications to cell-based therapies.

The baseline model predicts that cytolytic molecule secretion is strongly dependent on the IFFL pSFK → pSHP ⊣ pX, where X is either Vav or PLCγ. Interestingly, this motif is not uncommon in biological networks^86–89^ given its significant role in controlling cell activation by regulating signal transduction. Indeed, occluding the above subnetwork disinhibits signal transduction, allowing the NK cell receptor to continue signaling to its downstream mediators and thereby generate a greater response. In fact, there are many examples of increased cell activation using pharmacological inhibitors of phosphatases^20,90–93^. Here, we observed a similar result by decreasing the catalytic rate constant that regulates the rate of SHP activation *in silico*. Still, given the complexity of the signaling network, the efficacy of inhibiting the phosphatase is not immediately obvious. Our global sensitivity analysis revealed the importance of the phosphatase and simulating the effects of reducing phosphatase activity confirmed its impact, demonstrating the utility of the model.

The synNotch signaling system is a powerful tool for engineering cells with novel capabilities. One advantage of this approach is that it grants the researcher the ability to program new pathways in cells using established methods in molecular biology; however, arriving at the optimal signaling circuit via experimentation alone can be cumbersome, expensive, and time consuming. In addition, it is often not clear if optimality was obtained. Excitingly, mathematical models of cell signaling systems can help address the above barriers. Namely, the question of optimality can be addressed when a model is applied to perform optimal control, given a set of controllers and an objective function. In the work presented here, the initial amounts of the plasmid encoding lncRNA for the phosphatase, the plasmid encoding cytolytic molecules GZMB and PRF1, and the synNotch receptor are analogous to controllers as they help steer the NK cell signaling network to secrete more cytolytic molecules. Once a solution is found to the optimization problem, we gain insight on how to optimally control NK cell degranulation. Our findings demonstrate that the optimal strategy for maximizing the secretion of the cytolytic molecules is dependent on the number of rounds of stimulation that the cell will experience. We found that the optimal plasmid amounts have a bang-bang characteristic, meaning they are either applied at maximal dose or not at all. This is a known characteristic for bounded controls that appear linearly in the objective function^75^. In comparison, the optimal initial value of the synNotch receptor is more tunable and generally increases proportionally with the number of rounds of stimulation. Given that the synNotch receptor competes with the endogenous receptor for ligands, the increase in its initial amount reflects how much of the signal should be shifted to cytolytic molecule production and SHP inhibition versus secretion.

Our modeling and analysis predict the optimal amounts of the controllers for specific rounds of stimulation. As it stands, however, it is not completely understood how many target cells an individual NK cell can kill in its lifespan; that is, how many times an NK cell would be stimulated. Prager, et. al.^3^ recently showed that a given NK cell *in vitro* can kill up to six HeLa cells when confined to a microfluidic device. Elsewhere, others have demonstrated that NK cells are capable of serial killing^3,27,94–96^, implying they are able to undergo multiple rounds of stimulation. It remains to be seen if these results hold *in vivo* as well. While there is uncertainty in the exact killing potential, our analysis predicts that the highest secretion of GZMB and PRF1 occurs when we expect that NK cells will undergo many rounds of stimulation. The optimal synthetic biology solution is to have the cytolytic molecule-coding plasmid given at the maximal level in combination with a synNotch receptor to generate NK cells that continuously secrete effector molecules, even up to almost 10 rounds of stimulation. Although the precise numerical values of the optimal sets are subject to vary in practice, the qualitative predictions from the model are particularly useful; that is, the effects of using the maximum (or minimum) values of the plasmids and synNotch. Also, given that the intracellular pool of signaling species can limit the amount of secreted cytolytic molecules, and that the production of each species may not be feasible in practice, the optimal strategies proposed here complement ongoing research aimed at increasing the population of NK cells or their proliferative capacity along with the strategies presented here. Taken together, these approaches may improve the efficacy of NK cell-based therapies.

We acknowledge that the model predictions are sensitive to the assumptions made on the model (see methods). We assume an increase in the secretion of the cytolytic molecules will increase the likelihood of target cell death, since these effector molecules are the mediators for apoptosis. Given the model was fit to data from Srpan, et. al.^27^, where the ligands were immobilized in each well of the 96-well plate, we assumed the total amount of ligand in the system was fixed: *L*(*t*) + *C*(*t*) + *pC*(*t*) = *L*(*t*_0_) = constant. This, however, need not be true *in vivo* where the ligands are subject to degradation, shedding and clearance. The assumptions on the synNotch signaling pathway (see Table 3) were implemented to simplify the optimal control analysis. Certainly, the extracellular domain of synNotch receptor does not need to have the same properties as the endogenous receptor; the binding and internalization kinetics will be different in practice. Furthermore, the affinity between the transcription factor and the plasmid can vary depending on what specific nucleotide sequences are used in the promoter site and the specific transcription factor. In the future, we can simulate the model where one plasmid has a higher affinity to the transcription factor than another plasmid, which may affect the optimal strategy. We also assumed that lncRNA can bind to both SHP and pSHP with equal affinity. Fortunately, these assumptions can be addressed by experimentation and parameter estimation to improve the precision of the model. Lastly, we acknowledge that the solution to the optimization problem is not only sensitive to the model but also to the objective function. In particular, the constant *ρ* determined how much emphasis we placed on minimizing the given amount of exogenous material versus maximizing the cumulative amount of secreted cytolytic molecules. In our analysis, we placed equal weight on both. Future research can address this issue by solving the optimization problem for a variety of desired outcomes (e.g., more emphasis on performance).

Despite the limitations, the simulated results provide insight into NK cell secretion of cytolytic molecules GZMB and PRF1 mediated by the CD16 and NKG2D signaling pathways. We trained and validated a mathematical model by estimating the posterior distribution of the model parameters using the Metropolis-Hastings algorithm. An information-theoretic sensitivity analysis was subsequently performed on the model, from which we identified the inhibition of SHP strongly influences the secretion of the cytolytic molecules. By incorporating a synNotch signaling system, we found that the optimal conditions for maximizing degranulation is dependent on the number of rounds of receptor stimulation: we found that SHP inhibition is not optimal in the long term, while the production of the cytolytic molecules should be maximally induced almost all the time. In conclusion, the current work provides actionable insight into engineering robust NK cells with applications to immunotherapies.

## ACKNOWLEDGEMENTS

We immensely appreciate the Finley research group for critical evaluation of this manuscript as well as improvements to the model code. This work was supported a Viterbi/Graduate School Merit Fellowship (to S.Z.M).

## SUPPORTING INFORMATION

**File S1.** Computational model files (.m files with model code and .mat files with parameter values).

**File S2.** List of model species, reactions and parameters; provided as .xlsx file.

**File S3.** Supplementary figures; provided as .pdf file.

## Notes

### Competing Interest Statement

The authors have declared no competing interest.

