## Supplementary material for "An Optimal Control Approach for Enhancing Natural Killer Cells’ Secretion of Cytolytic Molecules": File S1: Driver.m

```
%% Driver file for model simulations

close all;
clear;
clc;

% Load parameter set
load('parameters.mat');

% NP (number of parameters)
% NR (number of estimates of each parameter)
[NP,NR] = size(parameters);

% Time domain (60 minutes)
t = linspace(0,60,601);

% Control sets
load('CD16controllers.mat');
load('NKG2Dcontrollers.mat');

% lncRNA-coding plasmid
u_CD16(1) = CD16_control(1,1); % no control set
u_CD16(2) = CD16_control(1,2); % max control set
u_CD16(3) = CD16_control(1,3); % semi-max control set

u_NKG2D(1) = NKG2D_control(1,1); % no control set
u_NKG2D(2) = NKG2D_control(1,2); % max control set
u_NKG2D(3) = NKG2D_control(1,3); % semi-max control set

% cytolytic molecule-coding plasmid
v_CD16(1) = CD16_control(2,1); % no control set
v_CD16(2) = CD16_control(2,2); % max control set
v_CD16(3) = CD16_control(2,3); % semi-max control set

v_NKG2D(1) = NKG2D_control(2,1); % no control set
v_NKG2D(2) = NKG2D_control(2,2); % max control set
v_NKG2D(3) = NKG2D_control(2,3); % semi-max control set

% synNotch receptor
R_CD16(1) = CD16_control(3,1); % no control set
R_CD16(2) = CD16_control(3,2); % max control set
R_CD16(3) = CD16_control(3,3); % semi-max control set

R_NKG2D(1) = NKG2D_control(3,1); % no control set
R_NKG2D(2) = NKG2D_control(3,2); % max control set
R_NKG2D(3) = NKG2D_control(3,3); % semi-max control set

%% Options for ODE solver

% Options without synNotch
options1 = odeset('RelTol',1E-06,...
                    'AbsTol',1E-09,...
                    'NonNegative',1:40);
                
% Options with synNotch                
options2 = odeset('RelTol',1E-06,...
                    'AbsTol',1E-09,...
                    'NonNegative',1:47);


%% Initial conditions (in uM)

% 40 species x 3 pathways (CD16, 2B4, NKG2D, respectively)

% Without synNotch
initvalue = zeros(40,3);
initvalue(1,:) = 3.8262E+01; % CD16
initvalue(2,1) = 0.0690; % Ligand CD16
initvalue(5,:) = 3.5300E-01; % 2B4
initvalue(6,2) = 6.67E-02; % Ligand 2B4
initvalue(10,:) = 3.0260E-01; % NKG2D
initvalue(11,3) = 0.0424; % MICA
initvalue(14,:) = 1.9100E+01; % SAP
initvalue(15,:) = 1.0500E+01; % SFK
initvalue(17,:) = 9.2300E+00; % ZAP70
initvalue(19,:) = 2.0500E-01; % LAT
initvalue(21,:) = 9.5200E-01; % PLCg
initvalue(23,:) = 7.9600E-01; % Akt
initvalue(25,:) = 5.2700E+00; % SLP76
initvalue(29,:) = 1.5000E+00; % Vav
initvalue(31,:) = 9.3500E+00; % Erk
initvalue(33,:) = 1.1800E+01; % SHP
initvalue(35,:) = 3.7600E+00; % SHIP
initvalue(37,:) = 10.38961039; % GZMB intracellular
initvalue(39,:) = 46.71770647; % PRF1 intracellular

x0_CD16 = initvalue(:,1); % CD16 only
x0_NKG2D = initvalue(:,3); % NKG2D only

% 47 species x 3 pathways (CD16, 2B4, NKG2D, respectively)

% With synNotch
initvalue1 = zeros(47,3);
initvalue1(1,:) = 3.8262E+01; % CD16
initvalue1(2,1) = 0.0690; % Ligand CD16
initvalue1(5,:) = 3.5300E-01; % 2B4
initvalue1(6,2) = 6.67E-02; % Ligand 2B4
initvalue1(10,:) = 3.0260E-01; % NKG2D
initvalue1(11,3) = 0.0424; % MICA
initvalue1(14,:) = 1.9100E+01; % SAP
initvalue1(15,:) = 1.0500E+01; % SFK
initvalue1(17,:) = 9.2300E+00; % ZAP70
initvalue1(19,:) = 2.0500E-01; % LAT
initvalue1(21,:) = 9.5200E-01; % PLCg
initvalue1(23,:) = 7.9600E-01; % Akt
initvalue1(25,:) = 5.2700E+00; % SLP76
initvalue1(29,:) = 1.5000E+00; % Vav
initvalue1(31,:) = 9.3500E+00; % Erk
initvalue1(33,:) = 1.1800E+01; % SHP
initvalue1(35,:) = 3.7600E+00; % SHIP
initvalue1(37,:) = 10.38961039; % GZMB intracellular
initvalue1(39,:) = 46.71770647; % PRF1 intracellular

x0_CD16_synNotch = initvalue1(:,1); % CD16 only
x0_NKG2D_synNotch = initvalue1(:,3); % NKG2D only

% no control == 1
% max control == 2
% semi-max control == 3
set = 1;

% Set the synNotch initial value
x0_CD16_synNotch(44) = R_CD16(set);
x0_NKG2D_synNotch(44) = R_NKG2D(set);

% Set the plasmid amount
parameters_synNotch_CD16 = [parameters;...
                            repmat(u_CD16(set),[1,NR]);...
                            repmat(v_CD16(set),[1,NR])];
                        
parameters_synNotch_NKG2D = [parameters;...
                            repmat(u_NKG2D(set),[1,NR]);...
                            repmat(v_NKG2D(set),[1,NR])];


%% Solve ODEs

disp(['Simulating ODEs ...']);

parfor i = 1:NR
    [~,CD16_results(:,:,i)] = ode15s(@CD16,t,x0_CD16,options1,parameters(:,i));
    [~,NKG2D_results(:,:,i)] = ode15s(@NKG2D,t,x0_NKG2D,options1,parameters(:,i));
end

parfor i = 1:NR
    [~,CD16_synNotch_results(:,:,i)] = ode15s(@CD16_synNotch,t,x0_CD16_synNotch,options2,parameters_synNotch_CD16(:,i));
    [~,NKG2D_synNotch_results(:,:,i)] = ode15s(@NKG2D_synNotch,t,x0_NKG2D_synNotch,options2,parameters_synNotch_NKG2D(:,i));
end

disp(['Simulations finished!']);
```
